# The effect of 0.01% atropine on ocular axial elongation for myopia children: a meta-analysis

**DOI:** 10.1101/2021.08.17.456658

**Authors:** Yan Yu, Jiasu Liu

## Abstract

**Objectives:** This meta-analysis aimed to identify the therapeutic effect of 0.01% atropine with on ocular axial elongation for myopia children.

**Methods:** We searched PubMed, Cochrane Library, and CBM databases from inception to July 2021. Meta-analysis was conducted using STATA version 14.0 and Review Manager version 5.3 softwares. We calculated the weighted mean differences(WMD) to analyze the change of ocular axial length (AL) between orthokeratology combined with 0.01% atropine (OKA) and orthokeratology (OA) alone. The Cochran’ s Q-statistic and I^2^ test were used to evaluate potential heterogeneity between studies. To evaluate the influence of single studies on the overall estimate, a sensitivity analysis was performed. We also performed sub group and meta-regression analyses to investigate potential sources of heterogeneity. We conducted Begger’ s funnel plots and Egger’ s linear regression tests to investigate publication bias.

**Results:** Nine studies that met all inclusion criteria were included in this meta-analysis. A total of 191 children in OKA group and 196 children in OK group were assessed. The pooled summary WMD of AL change was -0.90(95%CI=-1.25∼-0.55) with statistical significance(t=-5.03, p<0.01), which indicated there was obvious difference between OKA and OK in myopic children. Subgroup analysis also showed that OKA treatment resulted in significantly less axial elongation compared to OK treatment alone according to SER. We found no evidence for publication bias.

**Conclusions:** Our meta-analysis indicates 0.01% atropine atropine is effective in slowing axial elongation in myopia children with orthokeratology.

## Introduction

Myopia causes blurry vision when looking at distant object, which has become a worldwide healthy issue especially in some estern Ascian area.[1] There are about 1.406 billion myopia patients in the world, accounting for 22.9% of the total population. It is estimated that there will be 4.758 billion myopia patients in the world by 2050, accounting for 49.8% of the total population.[2] Myopia brings not only the decline of vision, but also serious complications caused by high myopia that can lead to irreversible vision loss, such as glaucoma, cataract, retinal detachment, retinal atrophy and other eye diseases.[3] At present, the number of children with myopia is increasing rapidly, especially in recent decades. Children with myopia are showing an increasingly younger trend, thereby increase the risk of high myopia.[4] Therefore, it is urgent to explore appropriate treatment to control the progression of myopia in children.

Axial elongation is the main cause of myopia, therefore, controlling axial elongation is important to prevent high myopia.[5] Current measures for controlling the progression of myopia include wearing glasses, orthokeratology lens, low concentration atropine and behavioral intervention.[6]

Atropine, as a non-selective M receptor antagonist, has been proved to have a significant control effect on the development of myopia[7]. There already has meta-analysis proved that the effect of atropine in different concentrations on myopia.[8] At present, 0.01% atropine is considered to have good curative effect, less adverse reaction, more stable and less rebound after drug withdraw.[9] Orthokeratology has been proved an effective means to control the progression of myopia in adolescents[10]. The mechanism of controlling the progression of myopia is that the hydraulic pressure generated by the contact lens temporarily reshapes the cornea, aiming to correct the distant vision by changing the shape of the central cornea, and secondly, it makes the peripheral cornea steeper to make the image focus in front of the peripheral retina, reduce the refractive error, and then achieve the best corrected vision [11]. Orthokeratology has a significant effect on the control of myopia progression, and has been accepted by doctors and patients.[12] There are a small number of studies have shown that the combination of orthokeratology and atropine can enhance the effect of myopia control. [13-15] However, due to individual differences, research groups, drug concentrations, and research design differences, the safety and effectiveness of the combined treatment still need to be verified. Therefore, the present meta-analysis aimed at determining the effect of 0.01% atropine on ocular axial elongation for myopia children.

## Methods

### Literature search

We searched PubMed, Cochrane Library, and CBM databases from inception to July 2021. The following keywords and MeSH terms were used: [“orthokeratology”] and [“atropine”] and [“ myopia”]. We also performed a manual search to find other potential articles.

### Eligibility criteria

1. Type of study. This study included high quality randomized controlled trials, cohort studies and case-control studies.
2. Type of patients. The patients should be children who undergone myopia. We will not apply any restrictions of race, age, education background, and economic status.
3. Intervention and comparison. This study compared OKA with OK for myopia control.
4. Type of outcomes. The primary outcome was ocular axial elongation.

If the study did not meet all of these inclusion criteria, it was excluded. The most recent publication or the publication with the largest sample size was included when the authors published several studies using the same subjects.

### Data extraction

Relevant data were systematically extracted from all included studies by two researchers using a standardized form. The researchers collected the following data: the first author’ s surname, publication year, language of publication, study design, sample size, age, follow-up time, ocular axial length, instrument, SER and ocular axial elongation.

### Quality assessment

The quality of the primary studies was assessed using the Cochrane risk of bias tool [16] by two independent researchers and an additional investigator in the case of any conflicts. The risk of bias for each study was evaluated according to selection bias, performance bias, detection bias, attrition bias, reporting bias, and other sources of bias. Each of these biases were classified as high risk (score 0), low risk (score 2) and unclear risk of bias (score 1). The total risk of bias was calculated by a summation of all categories.

### Statistical analysis

Review Manager 5.3 (The Nordic Cochrane Centre,Copenhagen, Denmark) and STATA version 14.0 (Stata Corp, College Station, TX, USA) softwares were used for the meta-analysis. We calculated the weighted mean differences(WMD) with their 95%confidence intervals(CIs) to analyze the change of axial length between OKA and OA. The Cochran’ s Q-statistic and I^2^ test were used to evaluate potential heterogeneity between studies. If significant heterogeneity was detected(Q test P<0.05 or I^2^ test>50%), a random effects model or fixed effects model was used. To evaluate the influence of single studies on the overall estimate, a sensitivity analysis was performed. We also performed sub group and meta-regression analyses to investigate potential sources of heterogeneity. We conducted Begger’ s funnel plots and Egger’ s linear regression tests to investigate publication bias.

## Results

### Characteristics of included studies

Initially, the searched keywords identified 36 articles. We reviewed the titles and abstracts of all articles and excluded 12 articles; full texts and data integrity were also reviewed and 15 were further excluded. Finally, 9 studies that met all inclusion criteria were included in this meta-analysis [13-15,17-22]. Figure1 showed the selection process of eligible articles. A total of 191 children in OKA group and 196 children in OK group were assessed. We summarized the study characteristics in Table1. Figure 2 and 3 represented the risk of bias in the primary studies based on the Cochrane risk of bias tool.

**Table 1.** Baseline charecteristics of all included studies

| First author | Language | Sample size |  | Age(Years) | follow-up<br>time(month) | Ocular axial length |  | AL measuring<br>instrument | SER(D) | Axial elongation (mm) |  |
| --- | --- | --- | --- | --- | --- | --- | --- | --- | --- | --- | --- |
|  |  | control | trial |  |  | control | trial |  |  | control | trial |
| Tang WT [13] | Chinese | 41 | 43 | 11.05±2.13 | 12 | 24.71±0.37 | 24.69±0.34 | IOL Master | -4.92±1.21/-4.90±1.16 | 0.20±0.05 | 0.12±0.03 |
| Shi YH [14] | Chinese | 89 | 86 | 12.5±3.6 | 6 | 24.95±0.17 | 24.87±0.22 | AL-Scan | -3.34±0.78/-3.31±0.79 | 0.25±0.11 | 0.11±0.09 |
| Shi MH [15] | Chinese | 66 | 62 | 10.88±1.34/11.06±1.50 | 12 | 24.99±0.66 | 24.85±0.48 | IOL Master | -3.29±0.89/-3.27±0.82 | 0.22±0.08 | 0.12±0.07 |
| Luo Y [17] | Chinese | 60 | 60 | 10.77±1.91/10.02±2.35 | 12 | 25.24±0.75 | 25.26±0.76 | A-mode ultrasound | -2.95±1.25/-3.00±1.32 | 0.22±0.21 | 0.13±0.19 |
| Chen Z [18] | English | 36 | 37 | 8.9±1.4/8.8±1.2 | 12 | 24.26±0.90 | 24.49±0.95 | IOL Master | -2.38±1.10/-2.85±1.08 | 0.30±0.11 | 0.21±0.15 |
| Kinoshita N [19] | English | 35 | 38 | 10.37±1.65/10.33±1.59 | 24 | 24.86±0.81 | 24.69±0.58 | IOL Master | -2.72±1.31/-2.60±1.29 | 0.40±0.23 | 0.29±0.20 |
| Kinoshita N [20] | English | 20 | 20 | 10.40±1.86/10.87±1.38 | 12 | 24.95±0.92 | 24.73±0.58 | IOL Master | -2.95±1.43/-2.81±1.43 | 0.19±0.15 | 0.09±0.12 |
| Tan Q [21] | English | 30 | 29 | 9.0±1.2/9.0±1.2 | 12 | 24.59±0.80 | 24.50±0.61 | IOL Master | -2.84±0.96/-2.65±0.92 | 0.16±0.15 | 0.07±0.16 |
| Vincent SJ [22] | English | 28 | 25 | 9.1±1.1/8.9±1.2 | 6 | 24.44±0.84 | 24.38±0.62 | IOL Master | -2.58±0.91/-2.38±0.81 | 0.05±0.08 | 0.01±0.12 |

AL:axial length; SER:spherical equivalent refractive; D:degree.

**Fig. 1.**
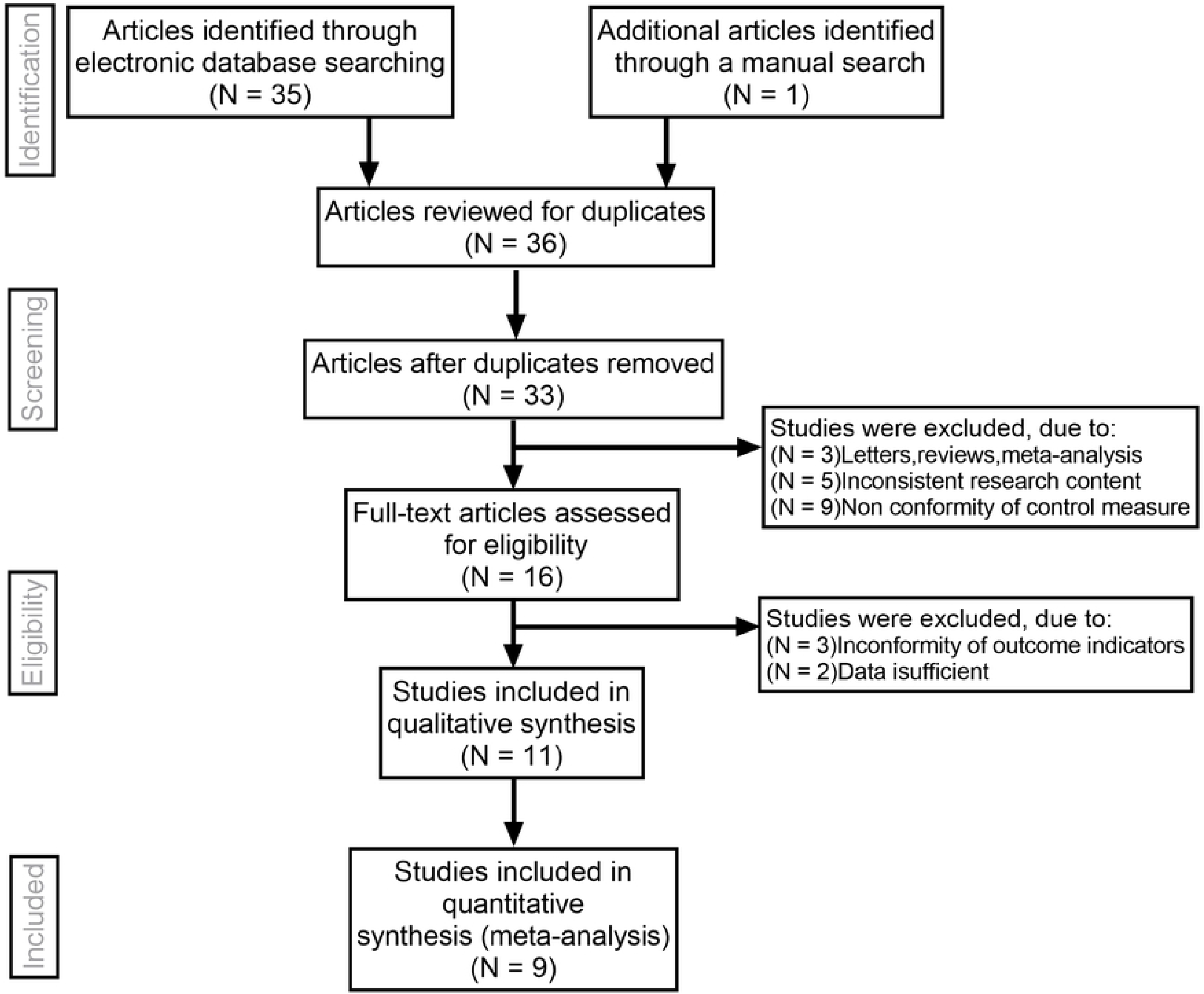
Flow chart of literature search and study selection. Nine studies were included in this meta-analysis.

**Fig. 2.**
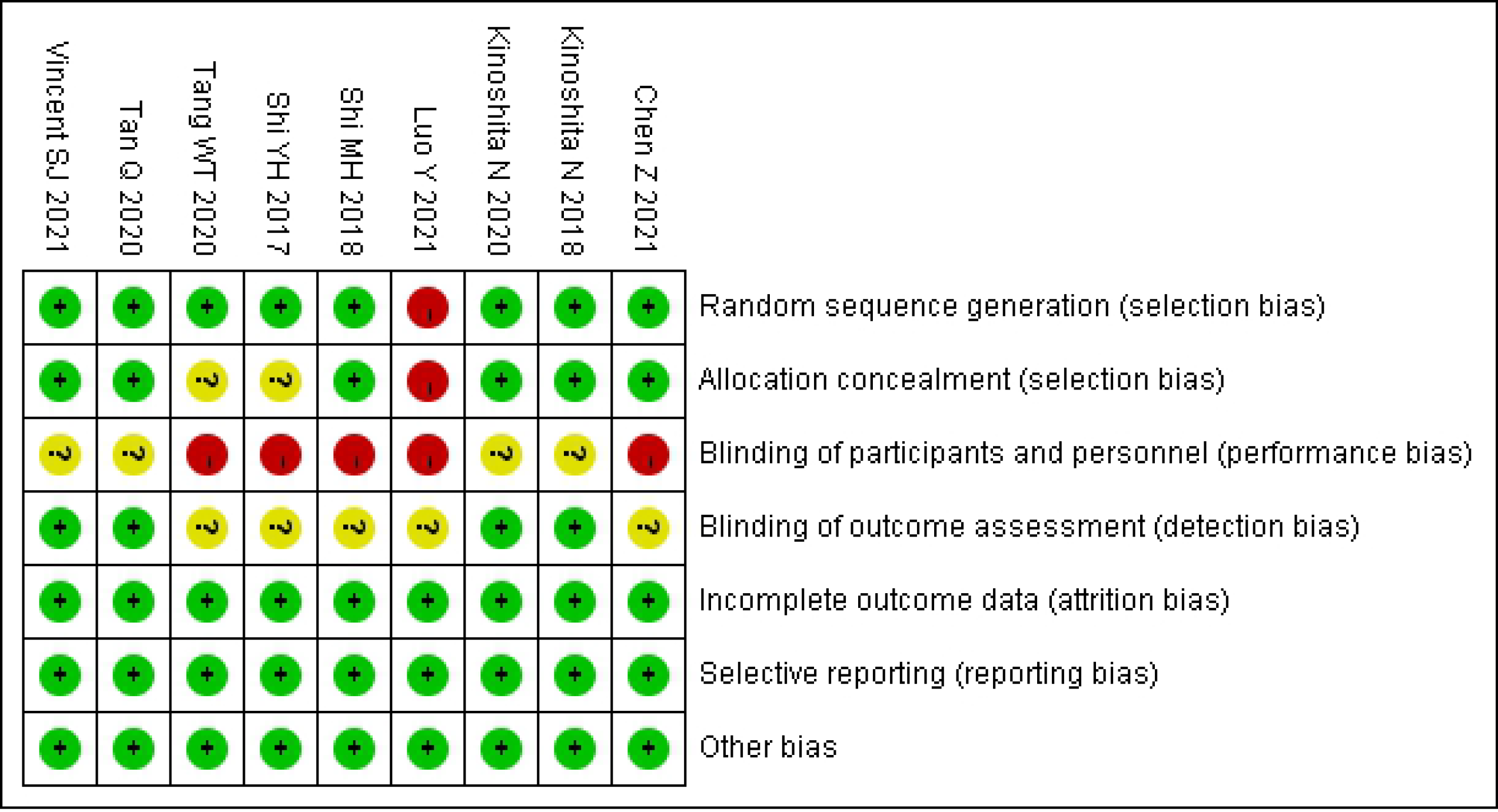
Risk of bias summary of quality evaluation of included randomized controlled trials.

**Fig. 3.**
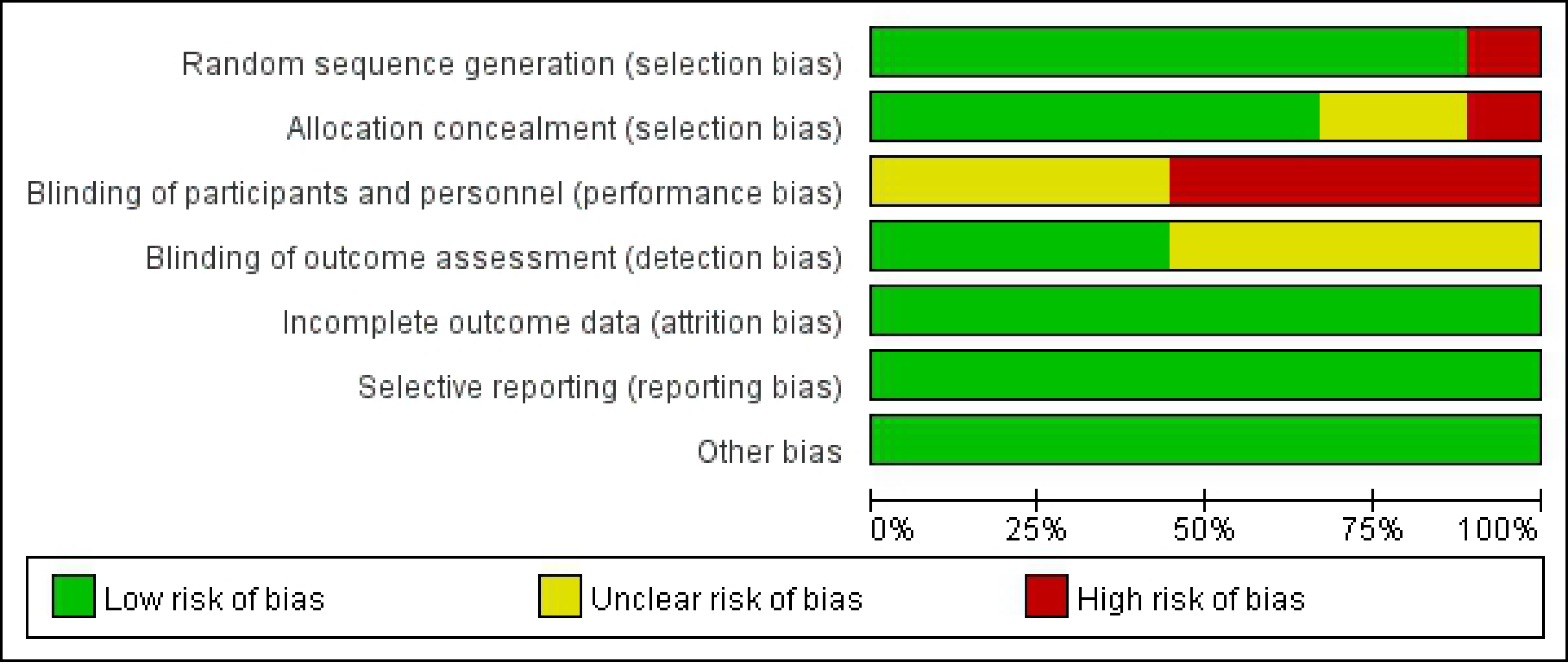
Risk of bias graph of quality evaluation of included randomized controlled trials.

### Quantitative data synthesis

The random effects model was used due to obvious heterogeneity among the studies(I^2^=81.4%, P<0.01). The pooled summary WMD of AL change was -0.90(95%CI=-1.25∼-0.55) with statistical significance(t=-5.03, p<0.01), which indicated there was obvious difference between OKA and OK in myopic children(Figure 4). Sensitivity analysis was carried out, and none of them caused obvious interference to the results of this meta-analysis(Figure 5). Meta-regression analysis conducted based on language, follow-up time, AL measuring instrument and SER. The result found that SER could explain potential sources of heterogeneity (Table 2). Subgroup analysis was performed according to whether SER was greater than 3. There was no significant heterogeneity within the group, but significant heterogeneity between the groups, which further explained that SER was the source of heterogeneity. The results of both groups showed that AOK treatment resulted in significantly less axial elongation compared to OK treatment alone(Figure 6). We found no evidence of obvious asymmetry in the Begger’ s funnel plots (Figure 7). Egger’ s test also did not display strong statistical evidence for publication bias.

**Fig. 4.**
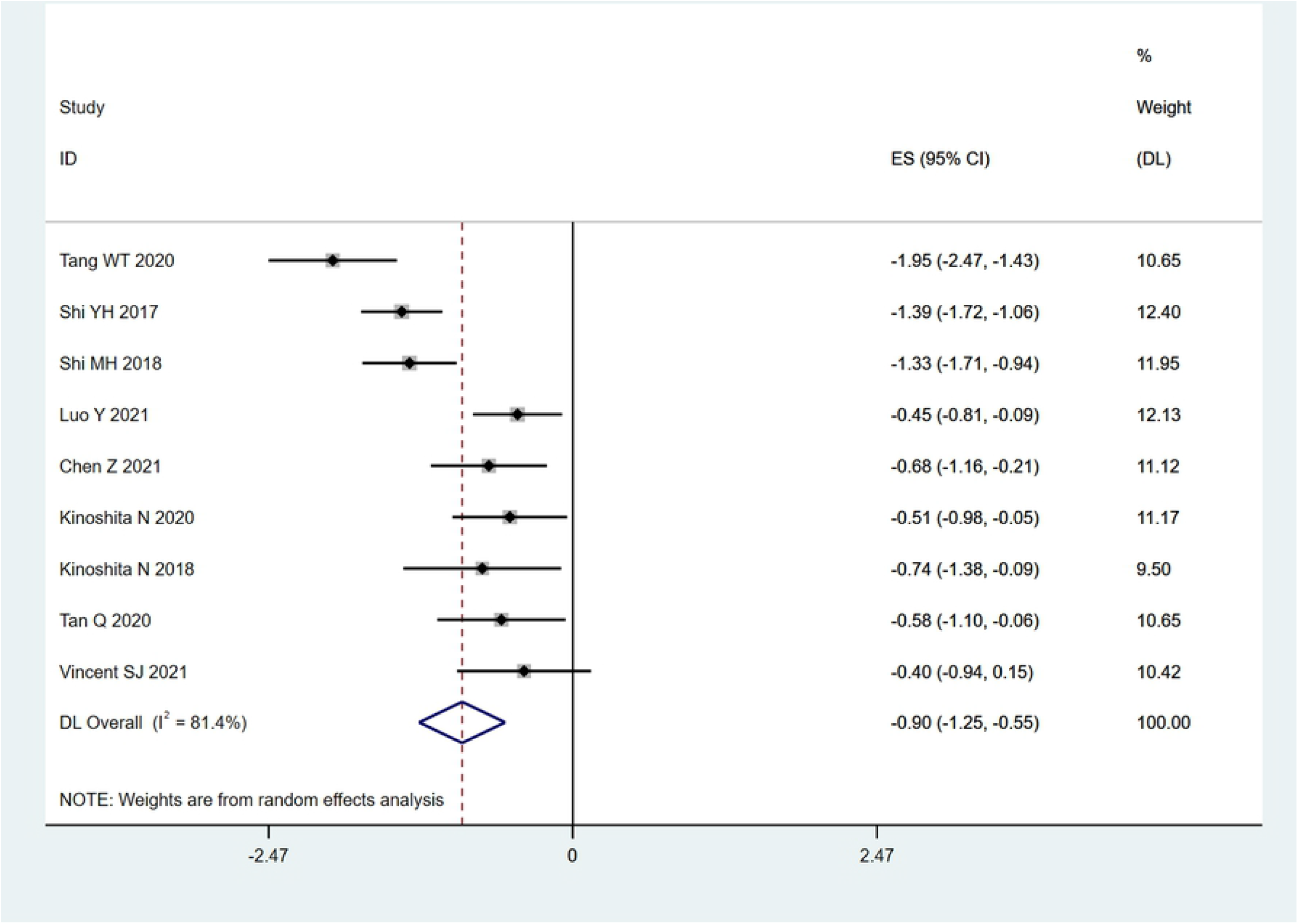
Forest plot of the effect of 0.01% atropine on ocular axial elongation for myopia children.

**Fig. 5.**
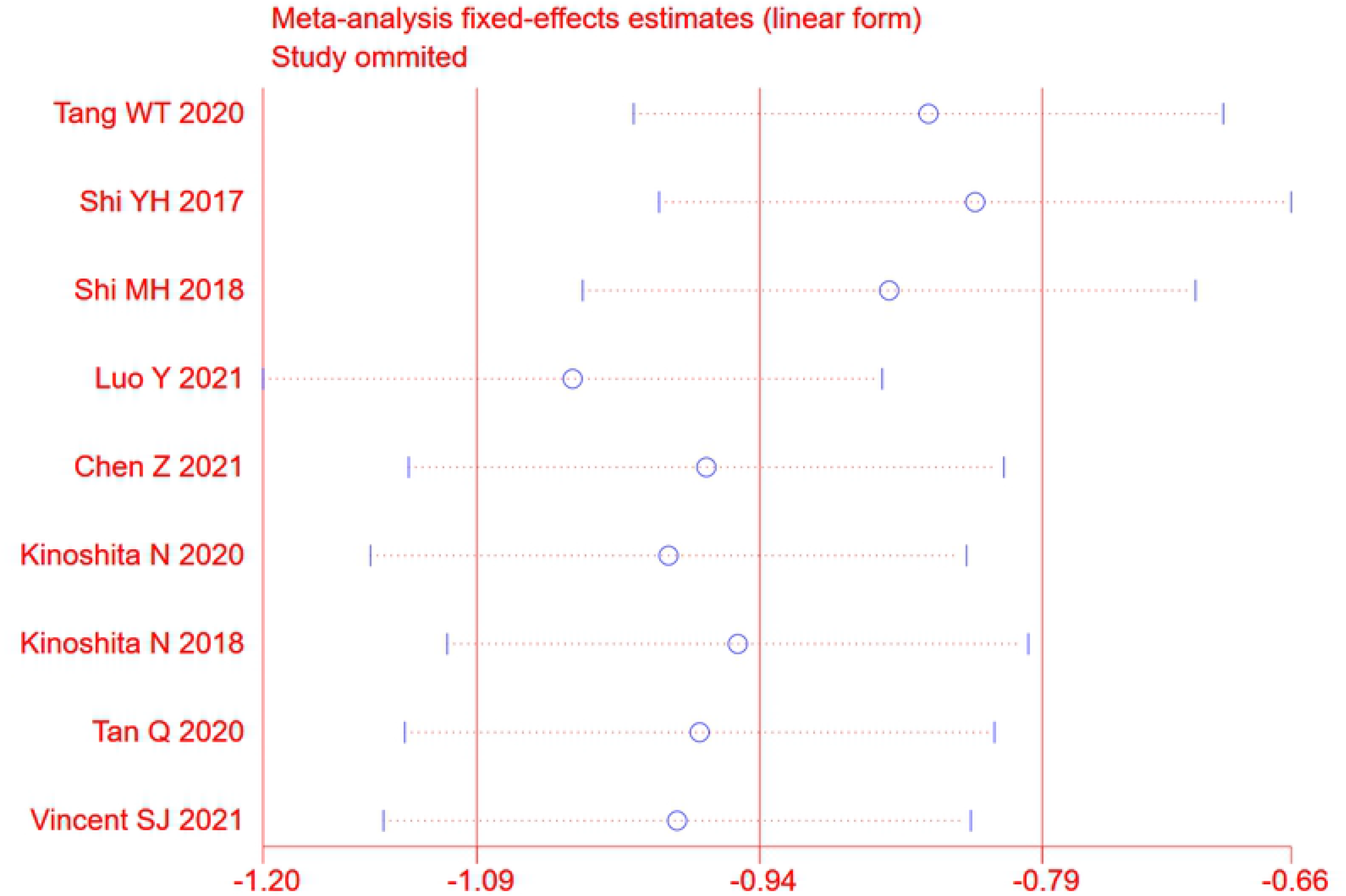
Sensitivity analysis. None of included studies caused obvious interference to the results of this meta-analysis.

**Table 2.**
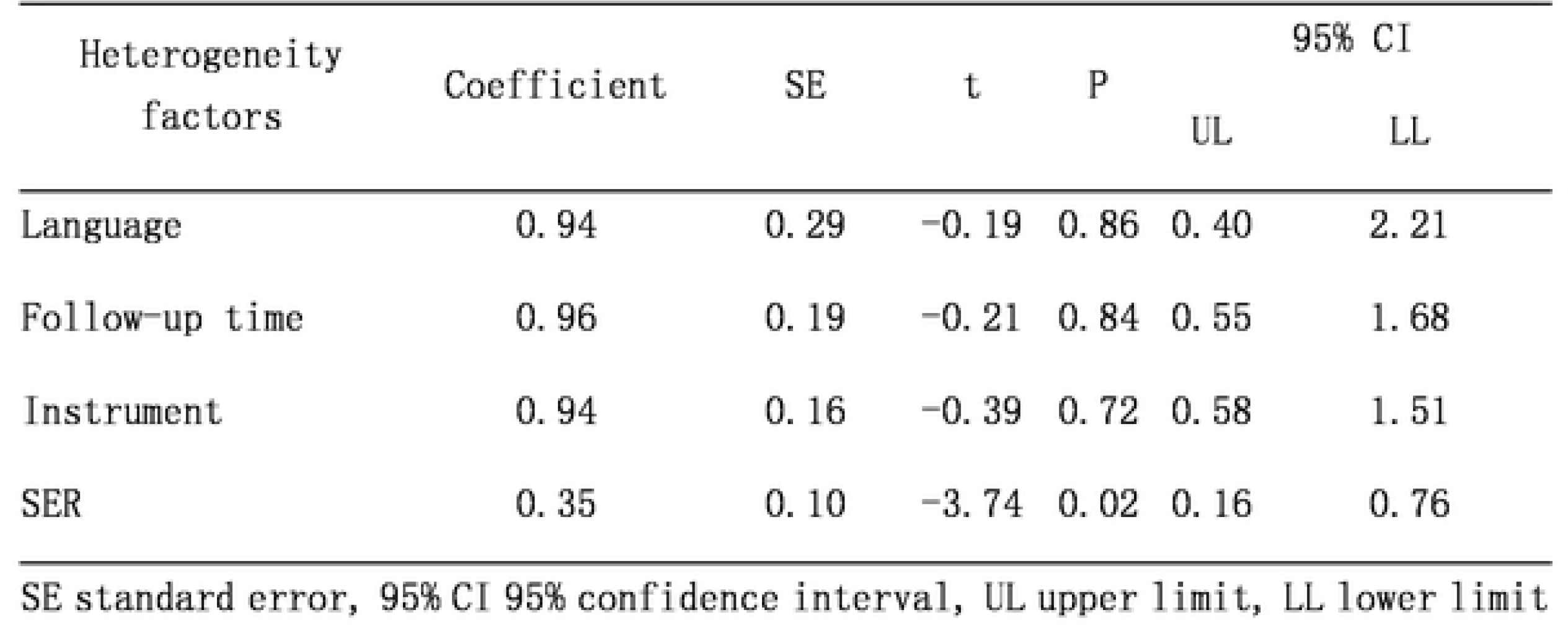
Meta-regression analyses of potential source of heterogeneity

**Fig. 6.**
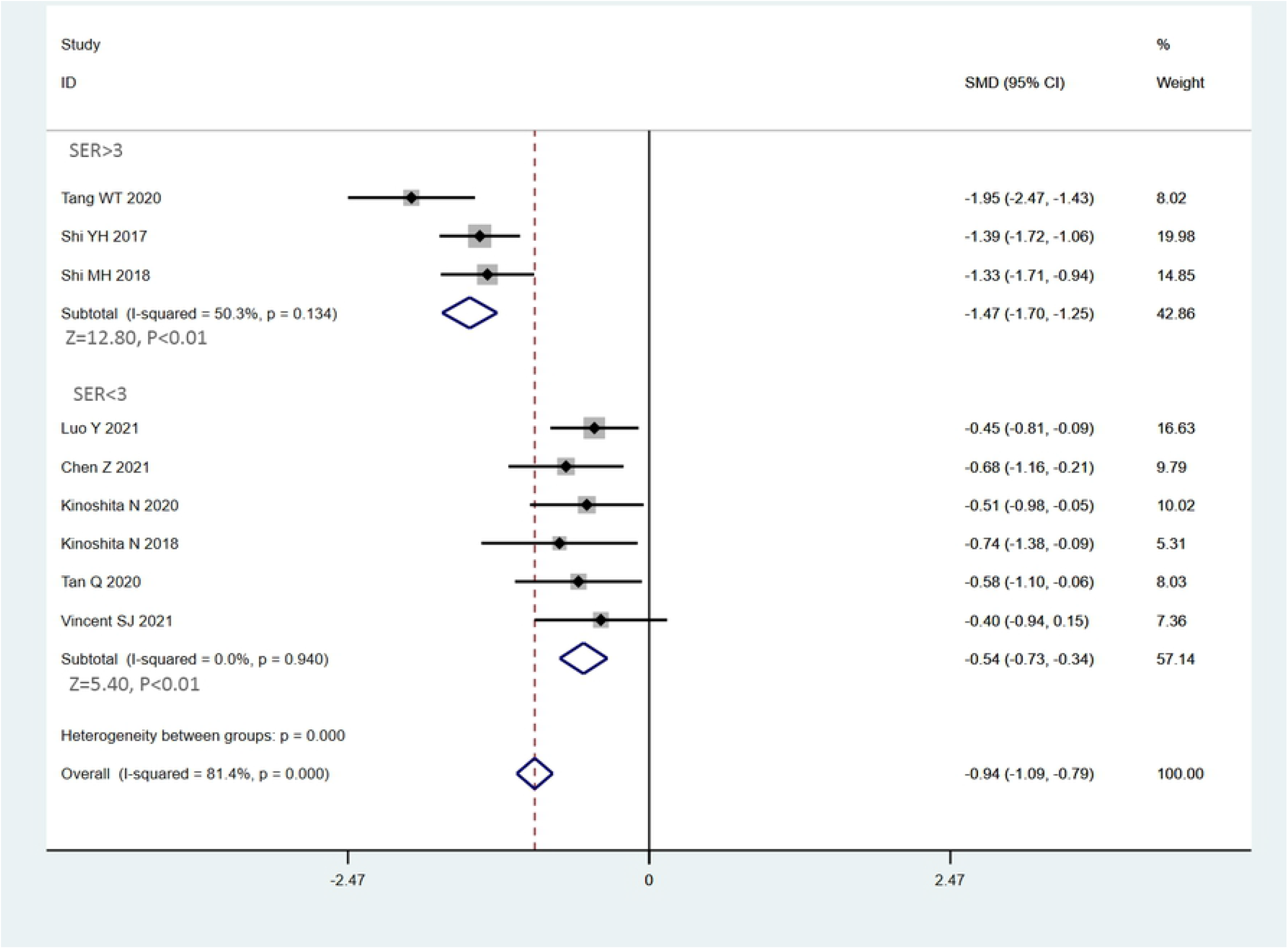
Subgroup analysis according to SER. Both groups showed AOK treatment resulted in significantly less axial elongation compared to OK treatment alone.

**Fig. 7.**
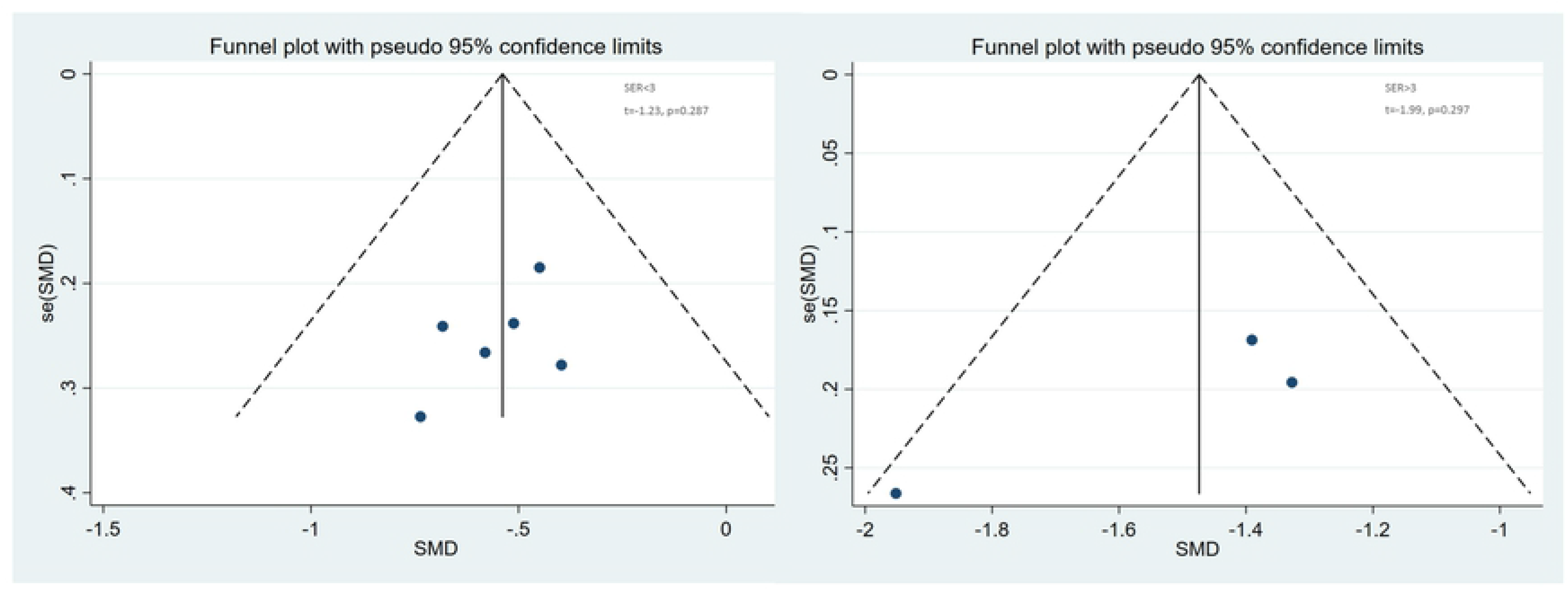
Begger’s funnel plot of publication bias. No publication bias was detected in this meta-analysis.

## Discussion

With the development of the information age, the incidence of myopia is increasing rapidly year by year, especially among yongner children[23]. The continuous progression of myopia will have a series negative effects[24]. As the loss of vision caused by high myopia is irreversible, myopia have become one of the main causes of untreated vision loss in the world[25]. The axis is an important monitoring index of myopia [21]. When the axis elongated beyond the normal value, axial myopia is occurred. There is a parallel relationship between the length of the axis and the progression curve of myopia corresponding to age.[26] Atropine is an alkaloid derived from belladonna and is a non-selective muscarinic acetylcholine receptor antagonist. Atropine eye drops act on the antimuscarinic receptors of the retinal, choroid and sclera. It may increase choroidal thickness by regulating dopamine release, or it may regulate scleral fibroblasts interferes with sclera remodeling in myopia. The effect of atropine suppressing myopia progression is dose dependent[27]. Some study believe that 0.01% atropine has a certain positive effect in controlling axial growth and suppressing myopia[20-22]. Orthokeratology lens adopts reverse geometric design and is made of highly oxygen-permeable materials. By changing the shape of the central cornea, the refractive power is changed. Studied from various mechanisms have confirmed that wearing an orthokeratology is one of the effective ways to control the progression of myopia [28]. Therefore, the efficacy of the OK combined with 0.01% atropine in the treatment of myopic children is a problem worth studying in particular. At present, there is a lack of multi center and large sample research in this aspect. This study aims to provide a comprehensive and reliable conclusion on the effect of 0.01% atropine combined with OK on ocular axial elongation for myopia children.

In the present meta-analysis, we systematically evaluated the effect of 0.01% atropine on ocular axial elongation for myopia children with OA. The 9 independent studies were included with a total of 191 children in OKA group and 196 children in OK group assessed. The pooled summary WMD of AL change was -0.90 with statistical significance, which indicated there was obvious difference between OKA and OK in myopic children. Because heterogeneity existed in the individual studies, subgroup analyses were conducted according to whether SER was greater than 3. Similar results were demonstrated in these subgroup analyses. The results of both subgroups showed that AOK treatment resulted in significantly less axial elongation compared to OK treatment alone. Furthermore, our results found no direct evidence of publication bias. Collectively, our findings strongly suggest that 0.01% atropine atropine is effective in slowing axial elongation in myopia children with orthokeratology , consistent with previous studies.

Still, our study has certain limitations. First, owing to the relatively small sample sizes and low level of quality of the included studies, there was insufficient data. Moreover, the retrospective nature of a meta-analysis can lead to subject selection bias. Importantly, the majority of included studies originated from Asia, which may adversely affect the reliability and validity of our results.

## Conclusion

Our meta-analysis suggests that 0.01% atropine atropine is effective in slowing axial elongation in myopia children with orthokeratology. However, due to the limitations, further detailed studies are required to confirm the present findings.

## Acknowledgments

We would like to acknowledge the helpful comments on this paper received from our reviewers. Also, we would like to thank all our colleagues working in the Ultrasound Department of the First Affiliated Hospital to Dalian Medical University.

